# Migration and Differentiation of Osteoclast Precursors under Gradient Fluid Shear Stress

**DOI:** 10.1101/379032

**Authors:** Y Gao, T. Y Li, Q. Sun, C. Y Ye, M. M. Guo, Z. B. Chen, J. Chen, B. Huo

## Abstract

The skeleton is able to adapt to mechanical loading through bone remodeling, i.e. bone resorption followed by bone formation. The osteoclasts close to microdamages are believed to initiate bone resorption, but whether local mechanical loading such as fluid flow regulates recruitment and differentiation of osteoclast precursors at the site of bone resorption has yet to be investigated. In the present study, finite element analysis first revealed that there exists low fluid shear stress (FSS) field inside microdamage. Basing on a custom-made device of cone-and-plate fluid chamber, finite element analysis and particle image velocimetry measurement were performed to verify the formation of gradient FSS flow field. Furthermore, the effects of gradient FSS on the migration, aggregation, and fusion of osteoclast precursors were observed. Results showed that osteoclast precursor RAW264.7 cells migrate along radial direction toward the region with lower FSS during exposure to gradient FSS stimulation for 40 min, obviously deviating from the direction of actual fluid flow indicated by fluorescent particles. When inhibiting calcium signaling pathway with gadolinium and thapsigargin, cell migration toward low-FSS region was significantly reduced. For other cell lines, MC3T3-E1, PDLF, rMSC and MDCK, gradient FSS stimulation did not lead to the low-FSS-inclined migration. After being cultured under gradient FSS stimulation for 6 days, the density of RAW264.7 cells and the ratio of TRAP-positive multinucleated osteoclasts in low-FSS region were significantly higher than those in high-FSS region. Therefore, osteoclast precursor cells may have special ability to sense FSS gradient and tend to actively migrate toward low-FSS region, which is regulated by calcium signaling pathway.

## Introduction

It has been well recognized that the skeleton is able to adapt to mechanical loading through bone remodeling, a process where mature bone tissue is removed followed by the formation of new bone tissue (1, 2). The removal of bone tissue at the early stage of bone remodeling is called bone resorption, which is mainly regulated by osteoclasts. Osteoclasts are multinucleated cells with the ability of dissolving bone mineral matrix. Mononuclear osteoclast precursors originate from hematopoietic stem cells and fuse into multinucleated mature osteoclasts under the activation of macrophage colony-stimulating factor (M-CSF) and Receptor Activator for Nuclear Factor-κ B Ligand (RANKL) (3). However, the recruitment or differentiation of osteoclast precursors at the site of bone resorption is still an unsolved problem.

Frost and his colleagues defined a specific anatomic structure of bone remodeling, basic multicellular unit (BMU), in which osteoclasts aggregate in and excavate the leading zone of BMU and osteoblasts deposit layers of osteoid in the reversal zone (1). It has been found that the resorption region characterized as BMU usually colocalizes with the microdamage in bone that forms after long-term cyclic mechanical loading (4, 5), which is called targeted bone remodeling (6). Further studies revealed that this colocation depends on the scale of microdamage. There are three types of microdamages in bone, i.e. linear microdamages, microdamages in a cross-hatched pattern, and diffuse damages (7, 8). The scale of linear or cross-hatched microdamages is approximately 10-100 μm, similar to or larger than that of osteoclasts (9); The size of diffuse damage is sub-micron, which is considerably less than that of osteoclasts. Some studies demonstrated that bone resorption usually occurs near large-scale microdamages (10-100 μm) rather than diffuse damage (10–12), but it remains unclear how osteoclast precursors migrate toward large-scale microdamages and further fuse into mature osteoclasts.

Previous studies have revealed that mechanical factors are included in targeted bone remodeling. For example, after inducing microdamage in the tibiae of 6-month-old rats, the number of resorption pits in weight-bearing animals was significantly higher than that in hindlimb-suspended group (13). This finding indicates that without the involvement of mechanical stimulations, chemical factors, such as the secretion from apoptotic osteocytes, are insufficient to cause bone remodeling around the microdamage. The deformation of bone mineral matrix caused by mechanical loading can drive fluid flow within cavities, which further induces fluid shear stress (FSS) on bone surface and adherent bone cells (14–16). Our previous studies have shown that osteoclast precursors migrate along flow direction and migration speed is proportional to the magnitude of FSS, which is regulated by calcium signaling pathway (9, 17, 18). The FSS level in a microdamage should be less than that on bone surface according to fluid mechanics (Fig. 1A), it therefore is hypothesized that osteoclast precursors may migrate toward the low-FSS region, such as around large-scale microdamages. The present study established the gradient FSS field by using a custom-made cone-and-plate flow chamber, through which the migration, aggregation, and differentiation of osteoclast precursors were observed and analyzed.

**Figure 1.**
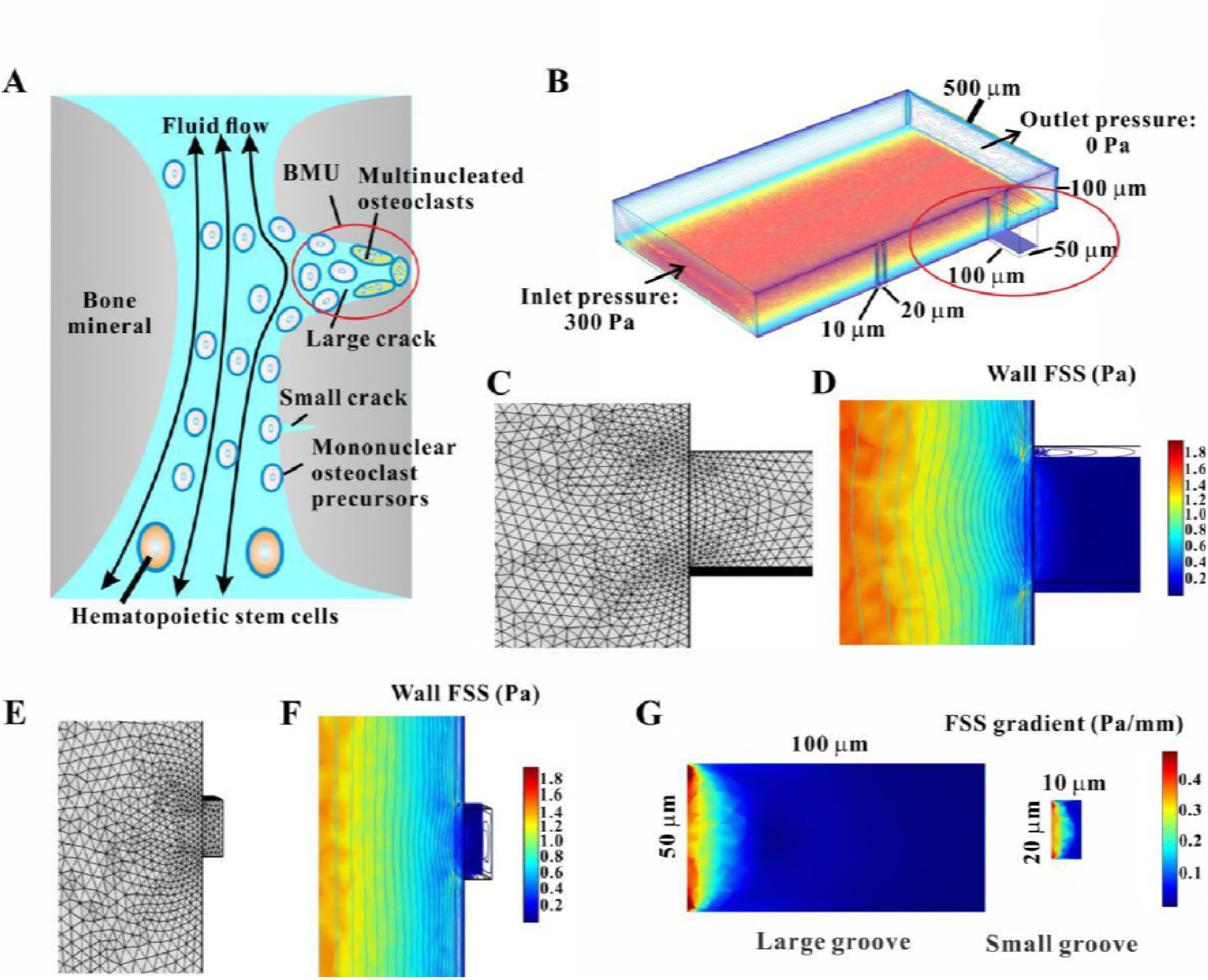
Fluid flow around microdamages in the cavities of trabecular bone. (A) Schematic of migration, aggregation, and fusion of osteoclast precursors around a large crack (red circle) regulated by FSS. (B) FE results of wall FSS for a fluid channel in which the grooves on the sidewall are constructed to mimic the microdamages with different sizes. The red circle indicates the region with low FSS around a large groove. FE mesh and wall FSS around the large groove are shown in (C) and (D), and those around the small groove in (E) and (F), respectively. (G) Wall FSS gradient on the bottom surface of the large and small grooves.

## Materials and Methods

### Cone-and-Plate Flow Chamber

We designed and fabricated a cone-and-plate flow chamber to establish a flow field with gradient wall FSS (Fig. 2A). This device mainly consists of motor, cone, and regular six-well culture plate. The distance between the cone’s tip and the plate surface was controlled by placing a silicon membrane with thickness of 0.4 mm on plate surface before fixing the cone and motor. The motorized rotation of the cone drives the medium in culture plate to flow over cells cultured on the plate. Wall FSS exerted on cells would be controlled by specifying the cone’s rotation speed.

**Figure 2.**
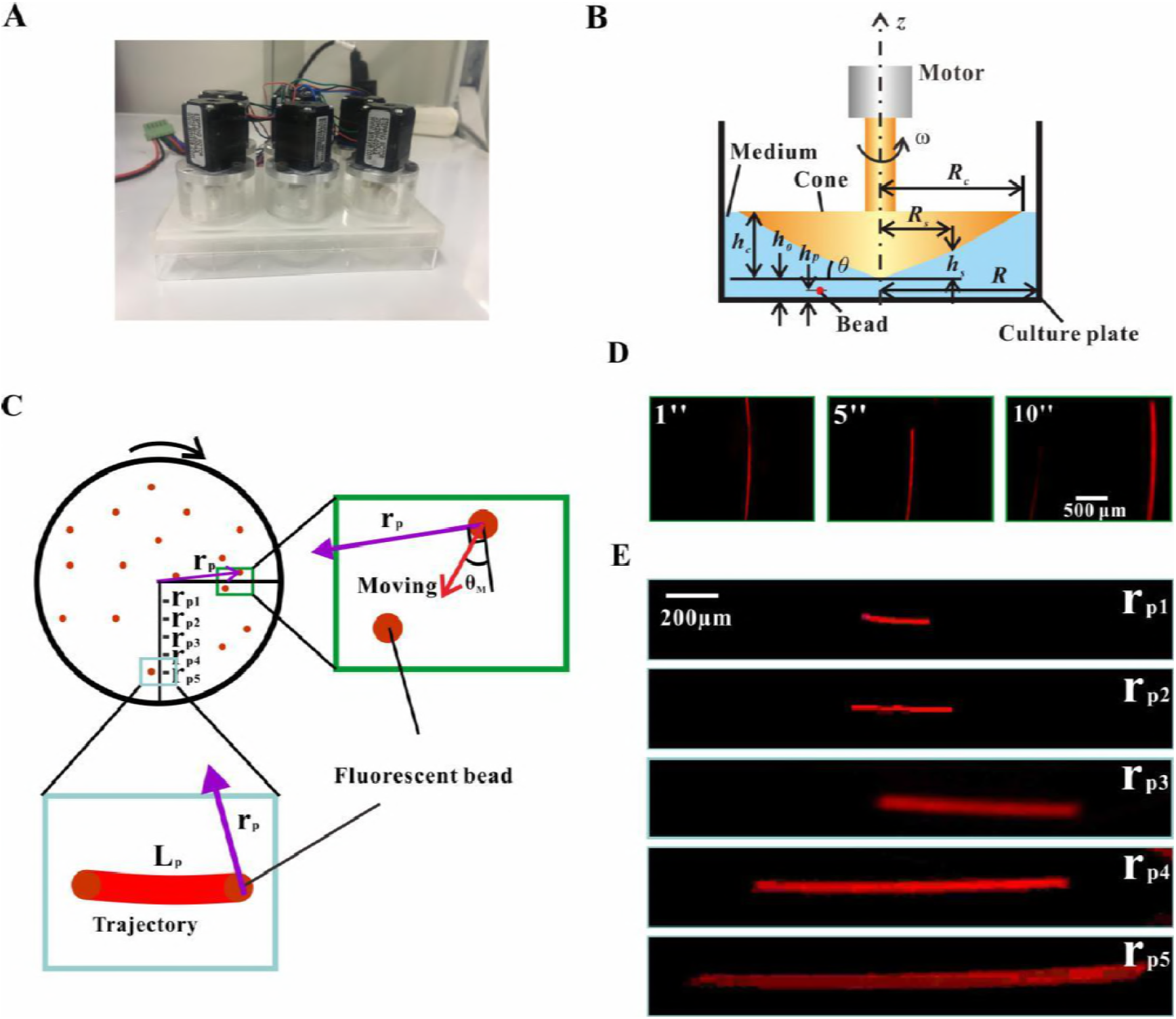
Cone-and-plate flow chamber and Particle Image Velocimetry. (A) Photo of the custom-made cone-and-plate flow chamber. (B) Diagram of cone-and-plate flow chamber, where *h*_0_ is the distance of the cone’s tip above the bottom of the plate, *h* is the height of the cone, *h_p_* is the height of beads in a specific focal plane. *θ* is the angle between the cone’s generatrix and the plate, *ω* is the angular velocity of the cone (18 rad/s), and *R* is the radius of the well. (C) Schematic of the analysis of PIV method, where the green and light blue boxes indicate the field of view in D and E, respectively. the orange dots are fluorescent beads, rp is the distance between fluorescent beads to the well’s center (r_p1_, r_p2_, r_p3_, r_p4_ and r_p5_ denote the locations of 2 mm, 4 mm, 6 mm, 8 mm and 10 mm away from the center, respectively), and *θ*_M_ is the angle between the actual flow directions (red arrow) indicated by beads’ movement and the circumferential directions. Red line means the beads’ trajectory. (D) Traces of fluorescent latex beads recorded at different times (6 mm from the center, time of exposure is 50 ms, *h_p_* is 0.2 mm). (E)Traces of fluorescent beads recorded at different location (time of exposure is 20 ms, *h_p_* is 0.1 mm).

### Numerical Simulation

Finite element (FE) analysis method was used for numerical simulation on flow field. An FE model was constructed considering the geometric and physical parameters of trabecular bone and microdamage (Fig. 1B). The length, width, and height of flow channel is 800, 500, and 100 μm, respectively, which is similar to the typical distance between the trabeculae (19). Furthermore, the grooves on the channel sidewall were set to mimic microdamages on the trabeculae. The depth and width of a large groove were 100 μm and 50 μm, respectively, and those of the small groove were 10 μm and 20 μm, respectively. The inlet and outlet pressures for this channel were assigned as 300 Pa and 0 Pa, respectively, on the basis of pressure difference in the lacunar-canalicular system (20). The mesh close to the grooves was locally refined to accurately simulate the wall FSS therein (Figs. 1C and 1D).

For the model of cone-and-plate flow chamber (Fig. 2B), the cone’s generatrix was machined as polyline to establish a wall FSS field with constant gradient on the plate surface. The radius of the polyline point *R_s_* was 8 mm, and its vertical distance *h_s_* to the cone’s tip was 0.05 mm. The total radius *R_c_* of the cone was 16 mm, and its vertical distance *h_c_* to the tip was 0.15 mm. The gap *h*_0_ between the cone’s tip and the plate surface was 0.4 mm. The angular velocity of the cone was 172 rpm. The radius *R* of a well for the six-well culture plate was 17 mm. No-slip boundary condition was assumed for all rigid surfaces in the model except for the cone, and a free-surface boundary condition was used for the upper fluid surface within the well.

COMSOL Multiphysics software was adopted for FE analysis. Incompressible viscous fluid in the above two models was assumed with density of 1×10^3^ kg/m^3^ and viscosity of 1×10^−3^ Pa·s. Digital image analysis was performed with ImageJ and MATLAB software.

### PIV Experiment

PIV technique was adopted to indicate the direction of fluid flow in the cone-and-plate flow chamber (Fig.2). Briefly, fluorescent beads of carboxylate-modified polystyrene with mean diameter of 0.5 μm (Sigma, USA) were suspended in deionized water (1:5) with a vortex mixer to ensure uniform distribution. After the cone-and-plate flow chamber ran stably for 5 min under fluorescence microscope, the movement of beads were recorded at 10 fps with a high-speed camera at the focal plane 0.2 mm above the plate, which were shown as arc-like trajectory in the images (Fig. 2D) (*Supplementary Movie S1*). The movement parameters of the beads, including speed and orientation, were then analyzed by an image-processing program based on MATLAB software.

PIV was also used to compute wall FSS close to the plate surface in the present study. The trajectory of beads’ movement was recorded at a focal plane 0.1 mm above the plate surface (Fig. 2C). The linear velocity *v*_p_ was obtained by dividing the length *L*_s_ of a trajectory arc with the exposure time of 0.1 s for one fluorescent image. Then wall FSS *τ_w_* = *η* · *v_p_* / *h_p_*, in which the distance *h_p_* is 0.1 mm and the viscosity is 1×10^−3^ Pa·s. This wall FSS value was compared with that from numerical simulation.

### Cell Culture

Osteoclast precursor RAW264.7 cells (European Collection of Cell Cultures, UK) were cultured in Dulbecco’s modified Eagle’s medium (DMEM, Hyclone, USA) supplemented with 10% fetal bovine serum (FBS, Gibco, USA), 100 unit/mL of penicillin (Sigma, USA), and 100 unit/mL of streptomycin (Sigma, USA) at 37 °C and 5% CO_2_. Osteoblast-like MC3T3-E1 cells and rat mesenchymal stem cells (rMSC) (American Type Culture Collection, USA) were cultured in alpha-modified Eagle’s medium (Hyclone, USA) supplemented with 10% FBS, 100 unit/mL of penicillin, and 100 unit/mL of streptomycin. The culture method for periodontal ligament fibroblasts (PDLF) and canine kidney epithelial cells (MDCK) (Peking Union Medical College Hospital, China) was the same as RAW264.7 cells. Mononuclear RAW264.7 cells were induced to fuse or differentiate into tartrate acid phosphatase (TRAP)-positive multinucleated osteoclasts by 100 ng/mL RANKL (Bio-Techne, USA) and 30 ng/mL M-CSF (Peprotech, USA). TRAP is a specific marker enzyme in osteoclasts.

### Time-Lapsed Imaging for Cell Migration

To observe cell migration under fluid flow, cytosolic calcium ions were stained with Oregon Green (Sigma, USA). The stock solution was prepared by mixing 50 μg Oregon Green with 18μL 10% Pluronic F-127 (Eugene, USA) in DMSO. Cells were seeded at density of 1×10^4^ cell/mL in one well of six-well plate and then incubated with the working solution, 2 μL stock solution of Oregon Green mixed with 1 mL DMEM medium, for 1 h at 37 °C with 5% CO_2_. The cells were washed three times with PBS and then immersed in 3 mL DMEM medium. After the plate was connected with a cone-and-plate device, live cell imaging was recorded for 40 min by 10 min intervals. Finally, migration trajectory of each cell was observed, and the migration parameters, including distance, speed and orientation, were analyzed.

### Inhibition of Calcium Signaling Pathways

To study the effect of calcium signaling pathways on the migration of RAW264.7 cells under gradient FSS, four pharmacological agents were employed. Similar to our previous studies (17, 18), the agents were incubated with cells before being exposed to FSS. In brief, 10 μM Gadolinium chloride (Gd, Sigma, USA) was supplied as the blocker of mechanosensitive cation channel (MSCC). The calcium stored in endoplasmic reticulum (ER) was depleted by 1 μM thapsigargin (TG, J&K Chemicals, China). 10 μM U-73122 (Darmstadt, Germany) was adopted to inhibit phospholipase C (PLC). For the above blocking tests, 10 min incubation of chemical reagents was applied. Calcium-free DMEM medium (Gibco, USA) was used to remove the extracellular calcium.

### Distribution and Differentiation of Cells under Long-Term Culture

RAW264.7 cells were subjected to FSS three times each day and once for an hour up to 6 days by using the cone-and-plate flow chamber. The chambers were placed in cell incubator and maintained at 37°C with 5% CO_2_. During this long-term culture, the cells at different distances to the center were counted each day. To observe the fusion of RAW264.7 cells, the cells on the culture plate were initially fixed with 4% paraformaldehyde (Wako Pure Chemical Industries, Japan) for 15 min and were stained with nuclear dye Hoechst (Eugene, USA) (PBS: Hoechst=1000:1) for 15 min at 37 °C after washing three times with PBS. TRAP Staining Kit was used to identify the formation of purple TRAP-positive, multinucleated osteoclasts.

### Statistical Analysis

Data were presented as mean ± SD. Statistical analysis was performed using a one-way analysis of variance with Tukey’s post hoc test for multiple comparisons to determine the statistical differences between the mean values of different groups. Each group in this study was repeated at least three times. A value of p<0.05 was considered statistically significant.

## Results

### Cone-and-Plate Flow Chamber Provides Flow Field Similar to Physiological Condition Close to Microdamage in Bone

An FE model, containing grooves with different sizes on the sidewall of a fluid channel, was established to investigate fluid flow field around microdamages in bone (Fig. 1B). Simulation results showed that the FSS level inside the large or small grooves is approximately 0.2 Pa, which is considerably lower than that of 1.6—1.8 Pa in the main flow path (Fig. 1D and 1F). It was further shown that the gradient of wall FSS on the bottom surface of large or small grooves ranges from 0 to 0.4 Pa/mm, and it is less than 0.1 Pa/mm in the most region inside the large groove (Fig. 1G).

Cone-and-plate flow chamber was designed in this study to establish a physiological gradient FSS field on the bottom of the plate. Figure 2E shows the images of fluorescent beads at different locations away from the center and it can be found that the lengths of the particles’ trace are positively related to the distance. The calculated wall FSS displayed a linear relation with the distance, ranging from 0.1 Pa to 0.7 Pa, and the fitted gradient is 0.6 Pa/mm (Fig. 3B). Numerical simulations also revealed low-FSS level in the center of culture plate (Fig. 3A) with the fitted gradient similar to PIV measurements (Fig. 3B). This FSS level and its gradient were selected to mimic the physiological FSS values of less than 3 Pa for bone cells (21) and to consider the above FE simulation result for large groove in Fig. 1G.

**Figure 3.**
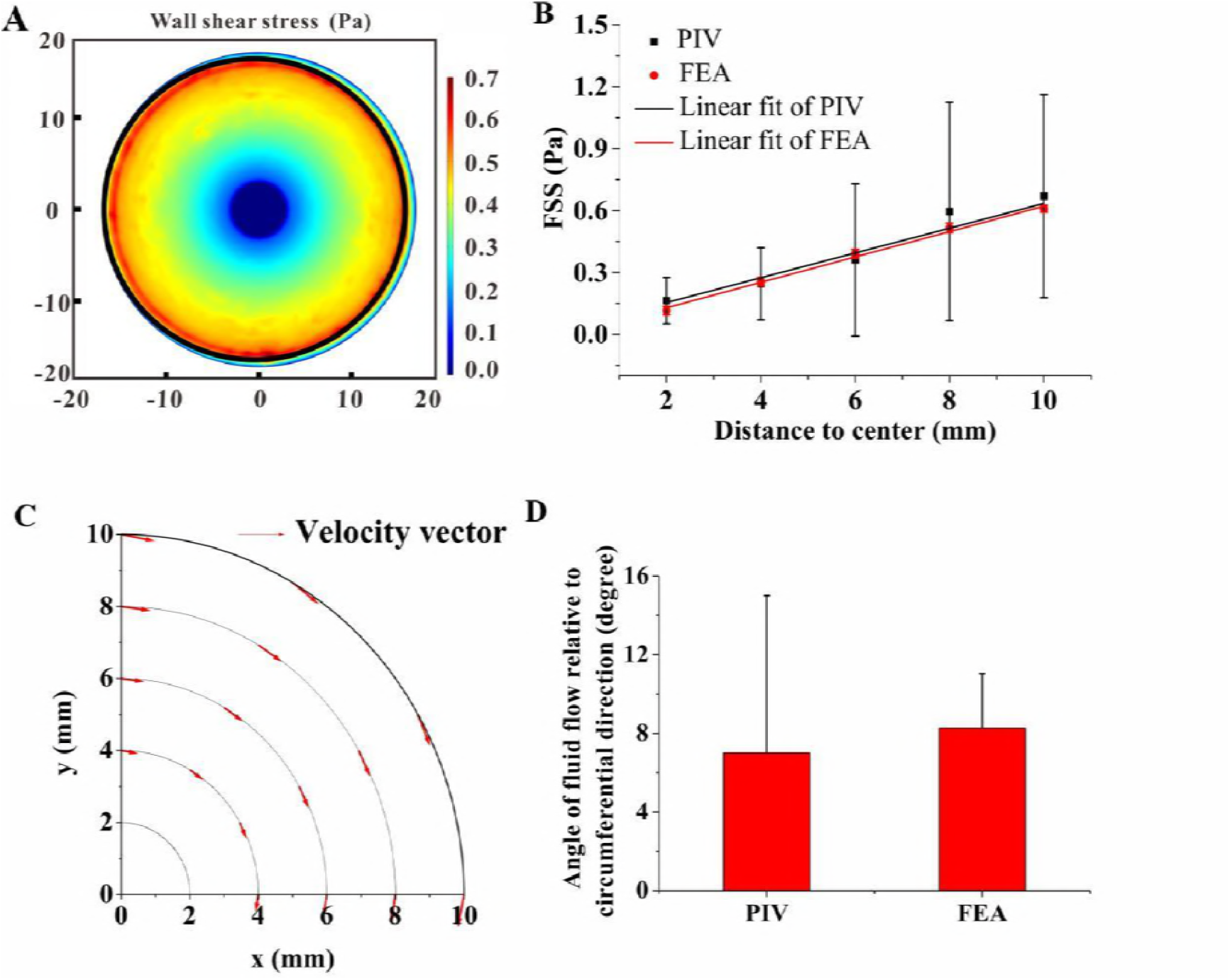
Finite element analysis (FEA) of fluid flow and comparation with PIV measurements. (A) Numerical simulation results of wall FSS on the surface of one well. (B) Wall FSS at different distance away from the center, in which the linear regression parameters for numerical simulation and PIV measurement are R=0.9979 and R=0.94553, respectively. (C) Velocity vectors (red arrows) indicating secondary flow against circumferential direction. (D) Angle of the direction of fluid flow relative to circumferential direction measured by PIV and predicted by FEA.

The velocity vectors of fluid flow at a specific point 10 μm away from the plate’s center are shown in Fig. 3C. It can be found that secondary flow deviating primary flow along circumferential direction was produced in the cone-and-plate flow chamber. PIV technique was adopted to experimentally indicate flow direction in the chamber (Figs. 2C and 2D). The average angles of flow direction relative to circumferential direction obtained from the FE analysis and PIV measurement were 8.3° and 7.0°, respectively, and no significant difference was observed between them (Fig. 3D).

### RAW264.7 Cells Migrate toward Low-FSS Region

After being stained with Oregon Green, the migration of RAW264.7 cells responding to gradient FSS was recorded for 40 min (Fig. 4A), and the images were captured at 6 mm away from the plate center (Fig. 4B; Supplementary Movies S2 and S3). The wall FSS at this region was approximately 0.4 Pa. The definition of migration parameters is shown in Fig. 4C. It can be found that the average angle of cell migration relative to circumferential direction after 40-min-FSS stimulation was approximately 90°, which was significantly larger than 69° or 75° after 10-min or 20-min FSS stimulation (Fig. 4D). No significant difference was observed for migration speed in all time periods, which ranges from 0.18 μm/min to 0.23 μm/min (Fig. 4E). The angle of cell migration relative to circumferential direction was considerably larger than that of secondary flow in Fig. 3D, suggesting that the migration of cells toward low-FSS region is not driven directly by fluid flow. When chemically blocking MSCC, PLC or ER, with Gd, U-73122 or TG, cell migration toward the plate center was significantly reduced compared with control group, which was the same as the removal of extracellular calcium ions with Ca^2+^-free medium (Fig. 4F; Supplementary Movies S4 to S7). For MSCC- and ER-blocking groups, cell migration along flow direction was enhanced (Fig. 4G), indicating that RAW264.7 cells lose the sensitivity for the gradient FSS.

**Figure 4.**
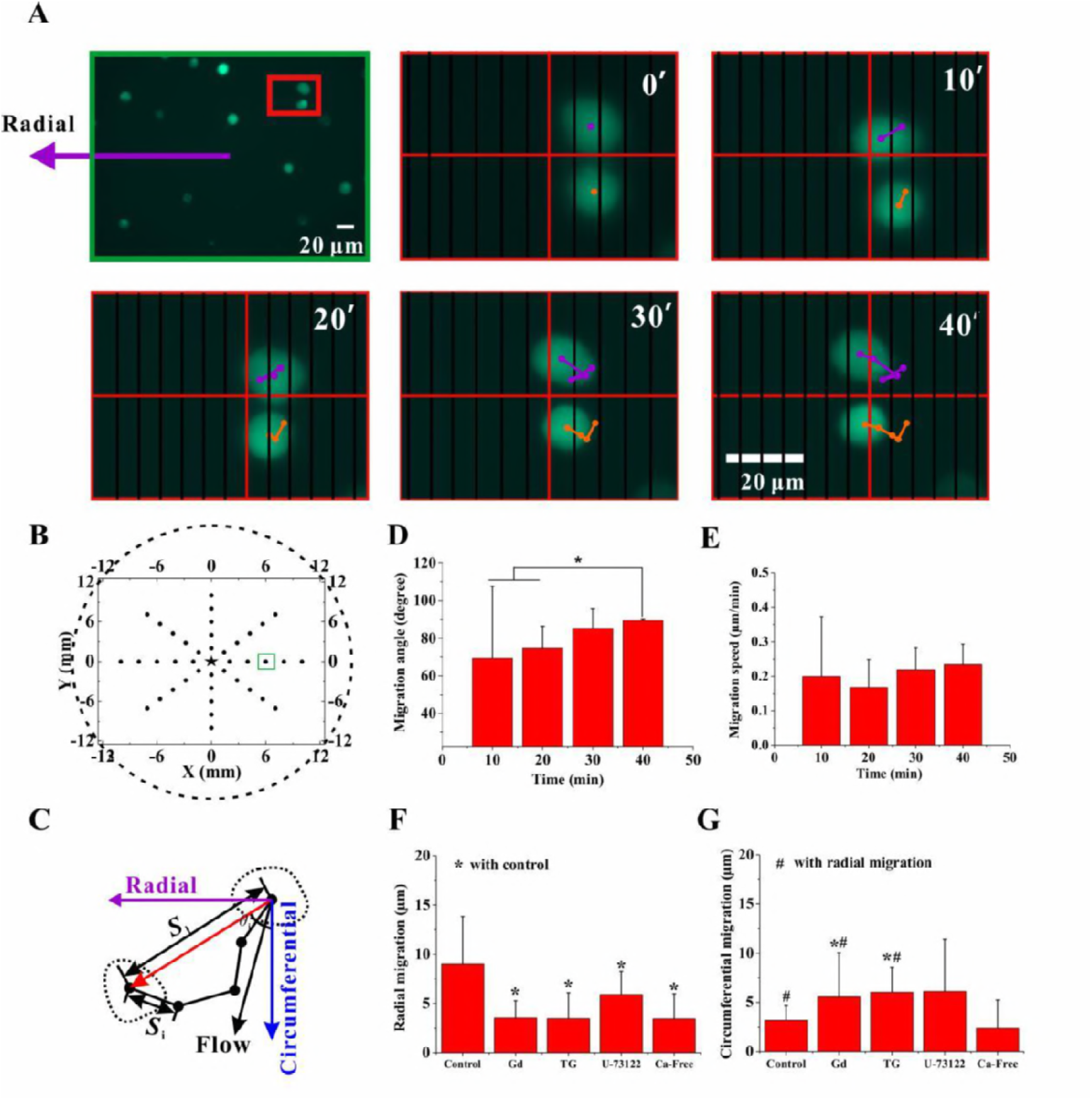
Cell migration under gradient FSS. (A) RAW264.7 cells stained with Oregon Green. The migration of the cells within a red box in the first image is shown in five time-lapsed images. The red and black lines are for the positioning of cell migration. (B) Positioning marks (black dots) in a well (dotted circle) of a six-well culture plate used for the multistage module of inverted fluorescence microscope. The black star marks the plate center. The green box indicates the field of view in A. (C) Definition of migration parameters. The black dots indicate the centroids of a migrating cell at different time. *S*_i_ is the distance of cell movement in a given time interval, *S*_1_ is the distance of a cell away from its initial position in a given time, and *θ*_i_ is the angle between circumferential direction and migration directions of one cell relative to its initial position. (D) Migration angle *θ*_i_ of cells in a given time interval. (E) Migration speed of cells in a given time interval. The effect of calcium signaling pathways on migration distance along radial (F) and circumferential (G) directions, respectively. *, #, p<0.05.

### Other Four Types of Cells Don’t Migrate along the Gradient of FSS

To examine whether gradient FSS-dependent cell migration is unique for osteoclast precursors or not, further experiments were performed for other four types of cells, i.e. MC3T3-E1, PDLF, rMSC and MDCK (Fig. 5A; Supplementary Movies S8 to S11). Under static condition, these cells did not reveal any tendency of migrating along radial or circumferential directions (Figs. 5B and 5C). After being exposed to fluid flow, only rMSC migrated toward the center within 40 min with significantly longer distance than MC3T3-E1, PDLF and MDCK, however, obviously less than RAW264.7 cells. The migration of RAW264.7, PDLF and rMSC along circumferential direction after flow stimulation was significantly enhanced compared with static condition, and the circumferential migration distance of PDLF and rMSC was significantly higher than RAW264.7 (Fig. 5C).The above results reveal that PDLF and rMSC tend to migrate along flow direction rather than radial direction, indicating they are not sensitive to gradient FSS.

**Figure 5.**
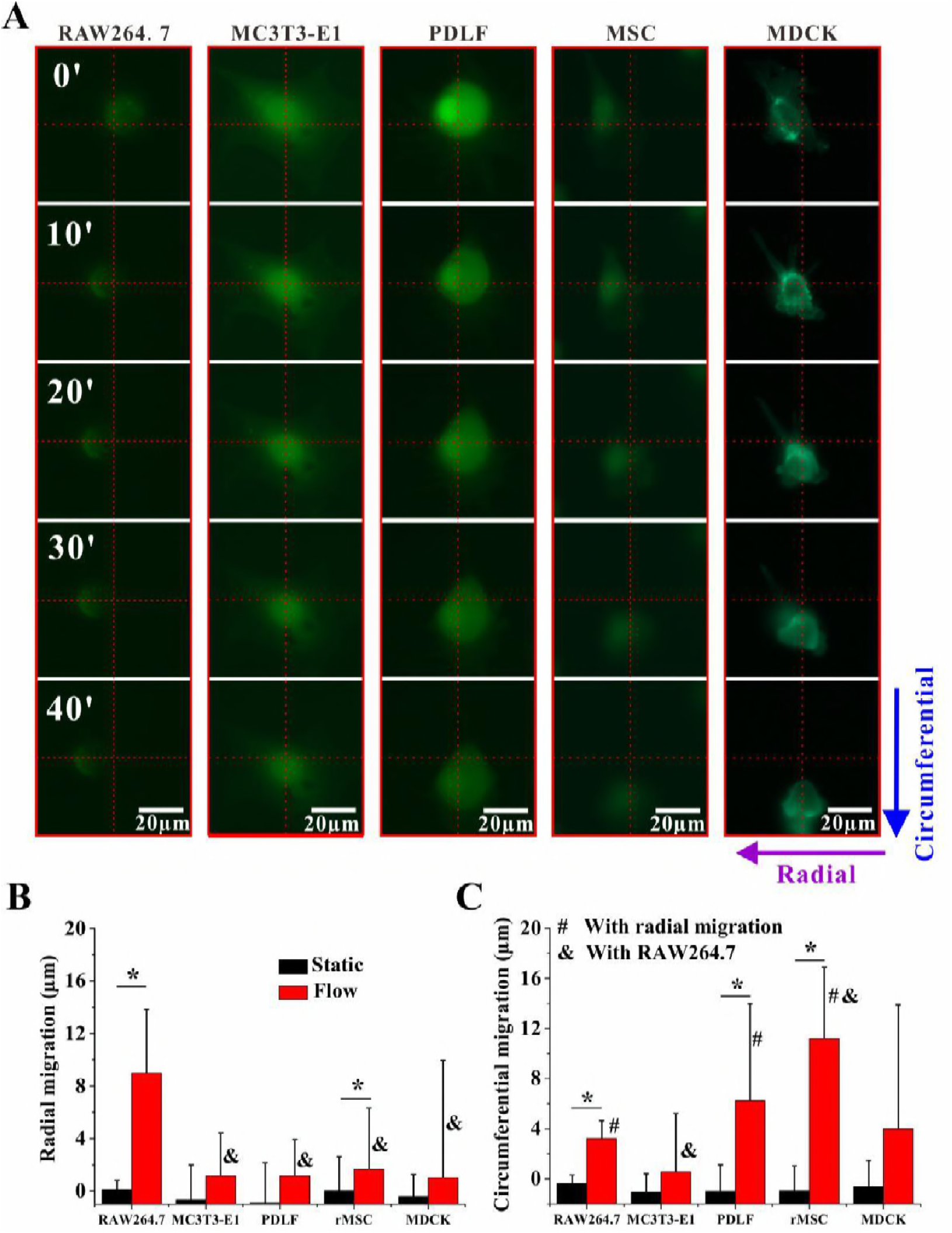
Migration of RAW264.7, MC3T3-E1, PDLF, rMSC and MDCK cells under gradient FSS. (A) The time-lapsed fluorescent images of migrating cells stained with Oregon Green. (B, C) The distance of cells migration along radial and circumferential directions, respectively. *, &, #, p<0.05.

### RAW264.7 Cells Tend to Aggregate and Differentiate in Low-FSS Region

To study the effect of gradient FSS-dependent migration on cell aggregation and differentiation, RAW264.7 cells were cultured in the cone-and-plate flow chamber system for 6 days and were exposed to flow stimulation for 1 h three times each day. The images at different distances relative to the plate center were captured at different times (Fig. 6A), and the number of cells was counted (Fig. 6B). When cells were cultured initially without fluid flow, no significant difference was observed for the cell number at different locations, approximately 104-120 cells in the field of view. At 3 days after flow stimulation, the cell number increased to about 160 at the center or 4 mm away from the center, which was significantly higher than 109 at the outer region of 8 mm to the center. After 6 days of culture and flow stimulation, the tendency of cell aggregation at the center became more evident and the cell number at the center was 364, which was considerably higher than 232 at the outer region. This higher level of cell density at the center may be due to cell migration toward the low-FSS region.

**Figure 6.**
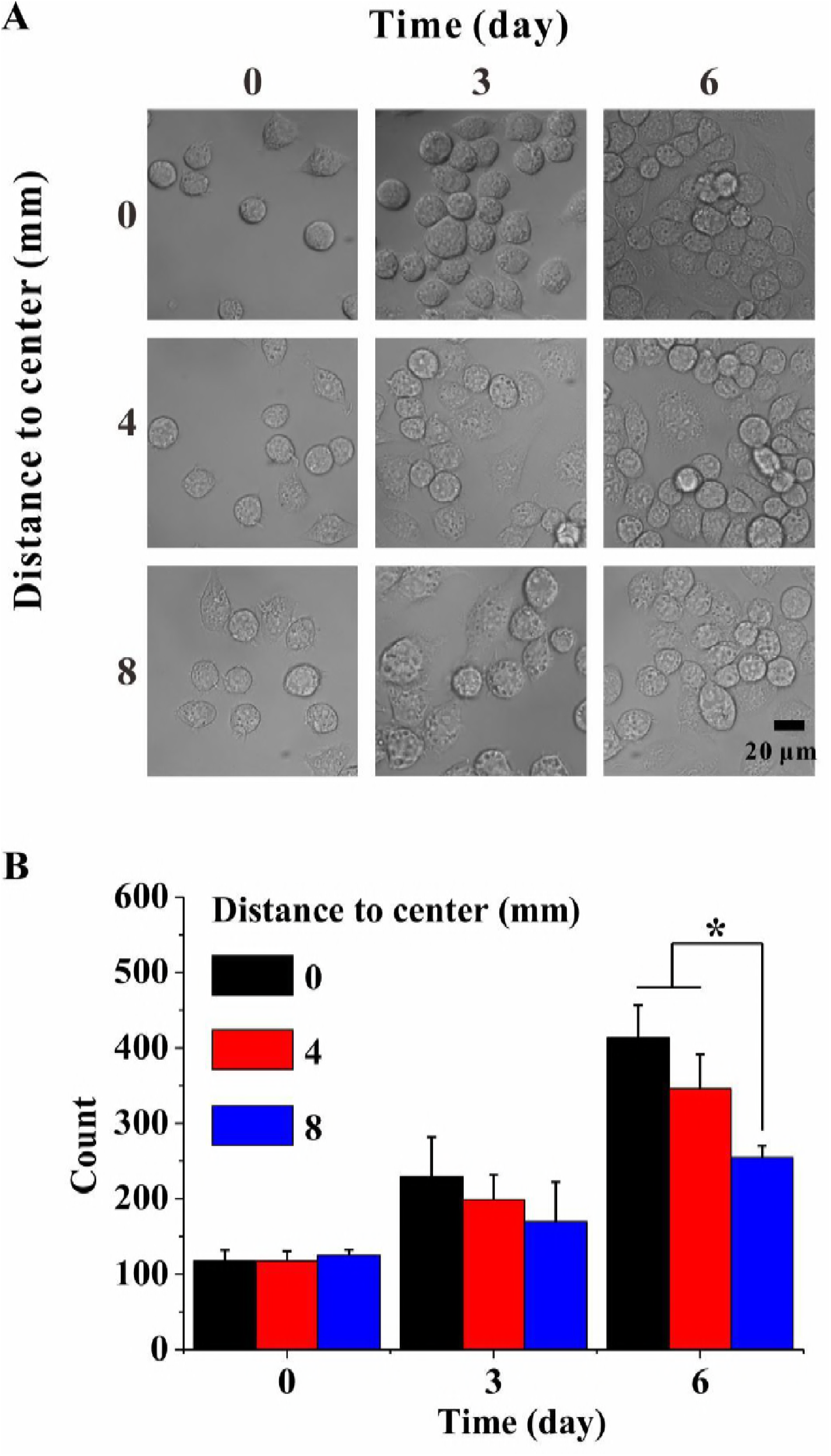
Cell aggregation under gradient FSS. (A) Photos of RAW264.7 cells at different distances to the center when being cultured for different times under 1-h fluid flow stimulation performed three times each day. (B) Average number of cells in the field of view. *, p<0.05.

To examine whether the fusion of RAW264.7 cells was also associated with FSS gradient or cell aggregation, the mature osteoclast marker TRAP and cell nuclei were stained and the images were taken at different locations and times (Figure 7A); the TRAP-positive cells with more than two nuclei were regarded as osteoclasts. Figure 7B shows the multinucleated osteoclasts at different distances of 0, 4 and 8 mm to the center before or 3 days and 6 days after flow stimulation. At the center of a well with a radius less than 4 mm, the percentage of multinucleated cells was highest up to 7.9%-8.8%, which was consistent with our previous work (9, 18). The ratio of fusion decreased with the increase of distance to the center (e.g., 3.3% at 8-mm region). Therefore, we could speculate that RAW264.7 cells were more likely to fuse into multinucleated osteoclasts in the low-FSS region.

**Figure 7.**
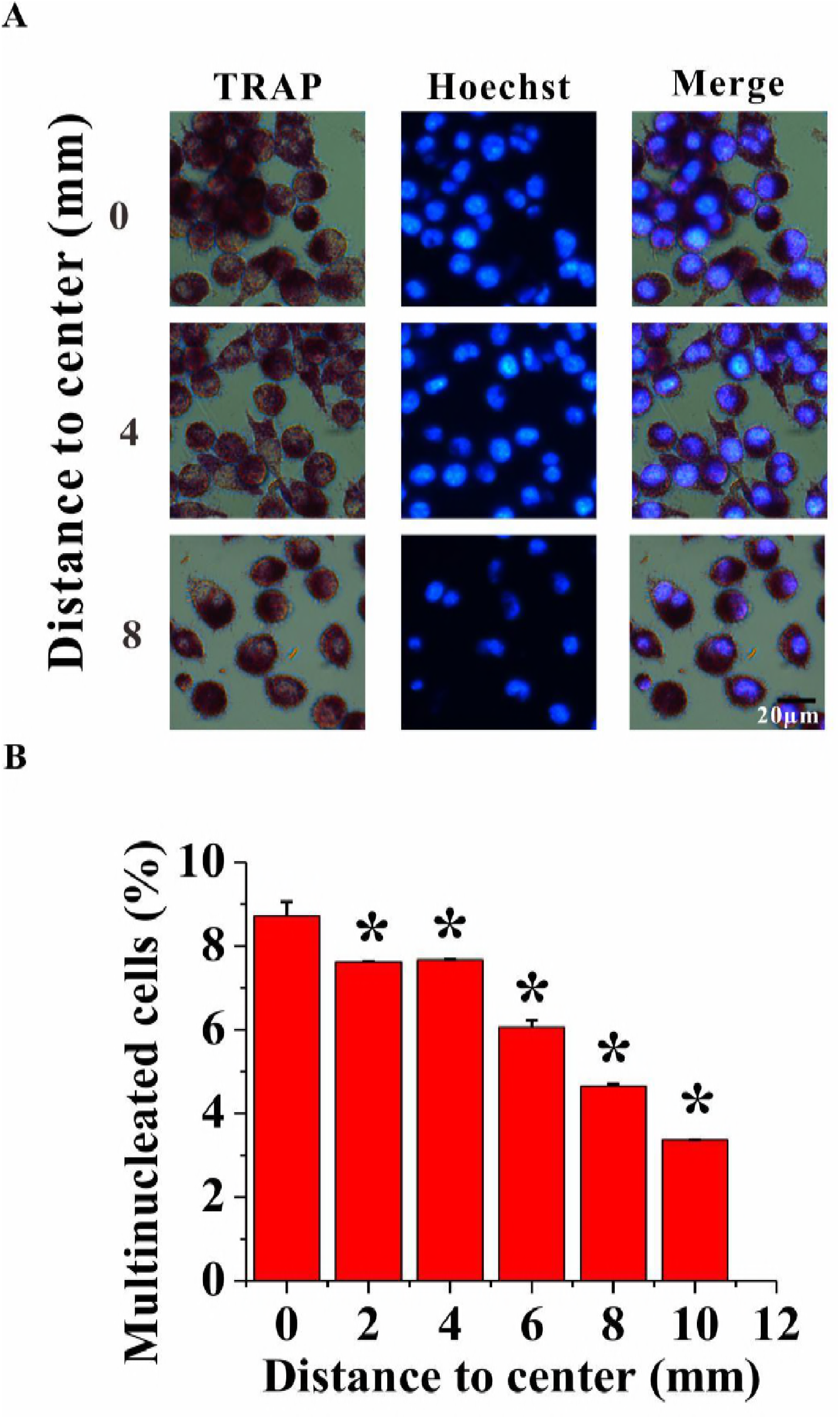
Cell differentiation under gradient FSS. RAW264.7 Cells were induced for differentiation by RANKL and M-CSF, and mechanically stimulated with gradient FSS up to 6 days, during which 1-h fluid flow stimulation was performed three times each day. (A) Images of RAW264.7 cells with nuclear staining by Hoechst and TRAP staining. (B) Percentage of multinucleated cells at different distances to the center. *, p<0.05.

## Discussion

In the present study, we constructed a gradient FSS field by using a custom-made cone-and-plate flow chamber system (Fig 2). Cone-plate configuration has been frequently used as viscometer in biomedical research, through which the rotation of a cone with linear generatrix may cause uniform FSS field (22). In the past several decades, cone-plate apparatus similar to cone-and-plate viscometer have been used to study uniform FSS on the adherent cells, however, only a few studies have been conducted about the effect of gradient FSS on biological behavior of cells (23–26). For example, Yan et al. adopted microfluidic technology to construct gradient FSS field and demonstrated that tumor cells prefers to adhere and grow at low-FSS region (i.e., the inner side of curved microvessels) (26). To the best of our knowledge, the gradient FSS field for the cone-and-plate system and its effect on cell migration have yet to be investigated.

In the early 1960s, Cox discovered that the direction of fluid flow in a cone-plate apparatus deviates circumferential direction, which is called as secondary flow (22). Some following theoretical analyses showed that at sufficiently low shear rate, the secondary flow is negligible and the primary flow along the circumferential direction provides a good approximation to the flow (23, 27). In addition, Sdougos et al. defined a non-dimensional Reynolds number Re = *ρωh*^2^ / (12*ν*), in which *ρ* is the density of medium, *ω* is the angular velocity, and vis the dynamic viscosity (28). The authors found that when Re was less than 0.5, the flow case was laminar flow; however, secondary flow would occur when Re was between 0.5 and 4.0. Recently, analytical and numerical analyses have described the steady, laminar, three-dimensional flow of a Newtonian fluid at low Reynolds numbers on endothelial cells (29). In the present study, Re was calculated as 0.06 considering that the height of the cone at the radius of 8 mm was 0.2 mm, and the flow case should be laminar flow. In fact, the cone-and-plate flow chamber designed in this study modified the cone’s shape; that is, the cone’s generatrix was not as linear as the traditional cone-plate apparatus adopted. This novel cone-plate flow chamber caused secondary flow, whose angle to circumferential direction was less than 8.3°, which was obtained by FE numerical simulation and PIV experimental test, respectively.

In the current work, RAW264.7 cells were exposed in a gradient FSS field ranging from 0.1-0.7 Pa (Fig 3), which was the physiological level within bone cavities (21). Surprisingly, cells did not migrate along the direction of fluid flow but almost move toward the low-FSS region under gradient FSS stimulation for 40 min (Fig. 4). And the directionality of cell migration was increased along with the time of flow stimulation (Supplementary Figure S12A). To our knowledge, the phenomenon of cells sensing to gradient FSS field and then migrating toward the low-FSS region have yet to be reported. When we exerted the gradient FSS on other four types of cells, i.e. MC3T3-E1, PDLF, rMSC and MDCK, none of them tended to migrate along radial direction (Fig. 5B and 5C). PDLF and rMSC even significantly migrated along circumferential direction compared with the radial component of their migration distance. Therefore, it seems that RAW264.7 osteoclast precursors have a special ability of sensing the direction of FSS gradient compared with other four types of cells.

We assume that this interesting phenomenon should be correlated with the opening of mechanosensitive ion channel under fluid flow. Our previous studies demonstrated that FSS induced more calcium responsive peaks in the late differentiated osteoclasts than the early ones, and MSCC, PLC and ER constituted the major signaling pathway (9, 18). We previously also found that when blocking the pathways of MSCC, ER, or extracellular calcium, the migration of RAW264.7 cells along flow direction is also significantly reduced (17). The present study showed that Gd and TG significantly inhibit the migration of RAW264.7 along radial direction and enhance the circumferential migration (Figs. 4F and 4G). And the directionality of cell migration was significantly reduced for MSCC-, ER- or PLC-blocking groups (Supplementary Figure S 12B). The above results indicate that calcium signaling pathways regulate the gradient-FSS induced-migration of RAW264.7 cells. But the molecular mechanism of this unique ability for RAW264.7 cells to sense gradient FSS is still unknown.

The process of osteoclast precursors migrating to the site of bone resorption and then fusing into mature multinucleated osteoclasts remains unclear. The apoptotic osteocytes near a microdamage releases cytokines, such as RANKL, phosphatidylserine, ICAM-3, or CD31, to regulate the recruitment and differentiation of osteoclast (30–33). Aside from the above chemical factors, some researchers have considered that mechanical strain in the bone matrix may regulate bone resorption. For example, an *in vivo* observation has shown that the local loading direction determines the tunneling orientation of osteoclasts (34). Through theoretical modeling and FE analysis, the strain around a BMU resorption cavity has been studied; the BMU has been predicted to move in the primary loading direction, and the osteoclast activity coincides with a low-strain region (35, 36). Our present study demonstrated for the first time that osteoclast precursors could sense the gradient of FSS and further migrate toward low-FSS region. In addition, the ability to sense FSS gradient is regulated by calcium channel. Finally, more monocytes fused into multinucleated osteoclasts at this region. This discovery may provide a new mechanism of osteoclast recruitment and differentiation during bone resorption and bone remodeling.

In the present study, the migration speed of cells was approximately 0.2 μm/min, which may lead to the average migration distance of 0.22 mm for 6-day culture under 1-h flow stimulation three times each day. The area of the outer section was larger than that of the inner section; thus, this inward cell migration caused more cells to aggregate toward the inner region. Notably, a higher FSS level in the outer region might cause more cells to be dead or detached from the substrate. However, we used physiological FSS and gentle stimulation strategy (i.e.,<1 Pa and 1-h stimulation three times each day); thus, abnormal adherent cells or cells suspended in the chamber were not observed during the 6-day culture (Supplementary Figure S13). However, when flow stimulation was applied all the time during 6 days, cell density significantly decreased after 1-dayculture, and considerably few cells were observed on the plate at 4 days (Supplementary Figure S14).

In conclusion, a custom-made cone-and-plate flow chamber was adopted in this study to construct a fluid flow field with gradient FSS. The experimental observation revealed that RAW264.7 cells did not migrate along the flow direction but directly moved toward the low-FSS region, which is regulated by calcium signaling pathways of MSCC and ER. This targeted migration led to the aggregation of osteoclast precursors at low-FSS region and finally enhanced their differentiation.

## Author contributions

Y. G. and B. H. designed the research; Y. G., B. H. and J. C. performed the research; Y. G., T. Y. L., C. Y. Y. and Z. B. C. performed numerical calculations; Y. G. performed the experiment; Y. G., T. Y. L., Q. S. and M. M. G. analyzed experimental data; Y. G. and B. H. wrote the article. All authors reviewed the manuscript.

## Acknowledgments

This work was supported by the National Natural Science Foundation of China [11572043 and 11372043 (BH)].

